# Volumetrically tracking retinal and choroidal structural changes in central serous chorioretinopathy

**DOI:** 10.1101/2023.09.18.557791

**Authors:** Bingjie Wang, Richard Brown, Jay Chhablani, Shaohua Pi

## Abstract

Central serous chorioretinopathy (CSCR) leads to accumulation of subretinal fluid and retinal thickness change, which can be readily detected in clinics using optical coherence tomography (OCT). However, current quantification methods usually require sophisticated processing such as retinal layer segmentations, and volumetric visualization of structural changes is generally challenging, which can hinder fast and accurate assessment of disease progression and/or treatment efficacy. In this study, we developed an algorithm that can register the OCT scans acquired from different visits without requiring prior layer segmentation and calculated the three-dimensional (3-D) structural change maps for patients with CSCR. Our results demonstrate that this tool can be useful in monitoring the progression of CSCR and revealing the resolution of pathologies following treatment automatically with minimal pre-processing.

## 1. Introduction

Central serous chorioretinopathy (CSCR) is a vision-threatening condition caused by the breakdown of the barrier function of retinal pigment epithelium (RPE) [1-3]. The hallmark of CSCR is the accumulation of fluid within the sensory retina and RPE, leading to serous retinal detachment [4-6]. Currently, observation is the standard of care for newly presenting CSCR. In most cases, acute CSCR is typically self-limiting, with retinal pigment epithelial detachment (PED) resolution and subretinal fluid (SRF) reabsorption spontaneously within three months [3, 7, 8]. However, up to 50% of acute cases can recur within one year [9], and 15% of acute cases may have persistent SRF longer than six months. These cases can result in chronic CSCR [10] that can cause permanent damage to photoreceptors [11] and may require treatment. Since there is no consensus on the best therapies for CSCR, long-term follow-up visits are critical to monitor disease progression and determine the appropriate course of action.

Optical coherence tomography (OCT) [12-14] has enabled the acquisition of high-resolution, three-dimensional (3D) retinal and choroidal images, and has been a routine imaging modality in the clinic. OCT has greatly improved the management of CSCR by allowing for the visualization of delayed resolution of SRF and abnormalities of RPE that are associated with a high risk of CSCR recurrence [15-17]. Furthermore, measurements of choroidal thickness obtained using OCT have been used to evaluate the progression of CSCR [18-20]. Additionally, OCT angiography (OCTA), which allows for the visualization of retinal and choroidal circulations, has been used to identify specific vascular impairments associated with CSCR [21-23]. However, these OCT/A evaluations are primarily performed in 2-D cross-sectional or *en face* images, which may not fully capture the 3-D nature of retina and pathologies [24]. Recently, 3-D quantification algorithms of OCT/A scans are emerging for better characterization of retinopathies in various diseases [24-26]. In a previous study, we developed a 3-D para-fovea vessel density (3D-PFVD) algorithm that achieved improved identification of foveal ischemia in diabetic retinopathy [27]. Based on these findings, we hypothesize that a 3-D quantification of the structure changes in CSCR might provide a better evaluation of the disease process and progression.

However, volumetric comparison of OCT/A scans acquired at different visits can be challenging due to mismatches resulted from the variation of pupil entrance for the scanning light beam (retinal curvature mismatch), shift of scanning regions (field mismatch), variation of transparency (reflectance mismatch), and eye movements (texture mismatch). All these can introduce inaccuracies in biomarker detection and measurement. To align two OCT/A volumes for comparison, registration is necessary. While we previously developed a volumetric registration algorithm for merging OCT/A scans from the same image session in rats [28], aligning and comparing OCT/A scans in humans and from different visits can be difficult due to substantial retinal structural deformation, especially for depth profiles in patients.

In this study, we proposed an improved automated volumetric registration algorithm that can align and compare OCT/A scans from baseline and follow up visits. Using this algorithm, we generated 3-D structural change maps between the scans to reveal the progression and resolution of pathologies in CSCR patients. We showed that this algorithm can detect subtle retinal and choroidal changes and therefore be a valuable tool for monitoring progression, guiding treatment decisions, and monitoring the effectiveness of treatments performed in CSCR patients.

## 2. Methods

### 2.1 Subjects

In this retrospective study, we included patients diagnosed with CSCR and compare with age-matched healthy subjects from available database from the UPMC Eye Center at the University of Pittsburgh, Pittsburgh, PA USA. The study was conducted following strict ethical guidelines, with all participants signing informed consent forms, in compliance with the requirements of the Declaration of Helsinki. Ethical clearance was obtained by the institutional review boards of the University of Pittsburgh. Eyes with significant ocular diseases, such as retinal vein or artery occlusion, were excluded. Participants with a history of intraocular surgery were also excluded, as these procedures may affect the OCT results and the interpretation of our findings. Additionally, images with a quality score below 6 or those exhibiting significant motion artifacts were excluded from the analysis.

In our study, we analyzed data from 10 eyes diagnosed with CSCR. The age of these subjects ranged from 26 to 60 years, with an average age of 50 ± 13 years. After their baseline scan, CSCR patients were advised to return to the clinic for follow-up scans with an average of 4 months (ranging from 2 to 9 months). The treatment group (n=5) primarily consisted of eyes that exhibited considerable vision challenges. Post the baseline evaluations, these patients underwent treatments such as laser photocoagulation (n=2 eyes), and photodynamic therapy (PDT) (n=3 eyes). The observational group (n=5) had eyes that retained a 20/20 visual acuity and remained untreated throughout the monitoring period.

### 2.2 OCT Image acquisition

The recruited subjects underwent OCTA imaging with a commercial swept-source OCT (SS-OCT) instrument (PLEX® Elite 9000, Carl Zeiss Meditec Inc., Dublin, CA). This device features a tunable laser with a center wavelength around 1050 nm and a bandwidth of 100 nm, allowing an A-line scanning speed of 100 kHz at a 3-mm tissue depth. The axial and lateral resolutions (optical) of the instrument were approximately 6.3-μm and 20-μm, respectively, and the axial resolution (digital) was approximately 1.95-μm. For our imaging, we centered OCT/A volumes on the fovea, covering a 12 mm × 12 mm field-of-view. This setup aimed to capture the majority of retina, including the fovea, optic disc, and peripheral regions. Each volume comprised 1024 A-lines per B-scan and 1024 B-scan locations per volume with two repeated B-scans acquired at each B-scan location. The complex optical microangiography (OMAG) algorithm was used to generate OCTA volumes [29]. To minimize potential motion-artifacts during imaging, FastTrac motion correction (Carl Zeiss Meditec Inc., Dublin, CA), which detects and tracks eye motion via line-scanning ophthalmoscope (LSO) fundus imaging, was employed. Further image processing of OCT/A was performed using the MATLAB R2022b platform (MathWorks, Natick, MA, USA), utilizing key functions such as “imgaussfilt”, “regionfill”, “detrend” and “circshift”, etc.

### 2.3 Volumetric registration

We developed a volumetric registration algorithm for the alignment of OCT/A volumes both laterally and axially. First, we identified the paired OCT/A volumes for the same patient from different visits (Fig. 1 A1,) and assigned the baseline scan as reference volume and the follow-up scan as moving volume. Second, we generated *en face* angiogram images (Fig. 1 B1) in both volumes by projecting first-15-maximum pixels of the OCTA signal along the entire A-lines. Then, we globally aligned the paired reference and moving *en face* angiogram images using cross-correlation, followed by local non-rigid registration of the moving image using a B-splines free-form deformation (FFD) model with the sum of squared differences (SSD) as similarity metric [30]. The calculated lateral transformation coefficient matrix could correct the mismatch between the vascular patterns (Fig. 1 B2) and was applied to all depth planes of the moving OCT volume as it was assumed to be the same across the entire depth.

**Fig. 1.**
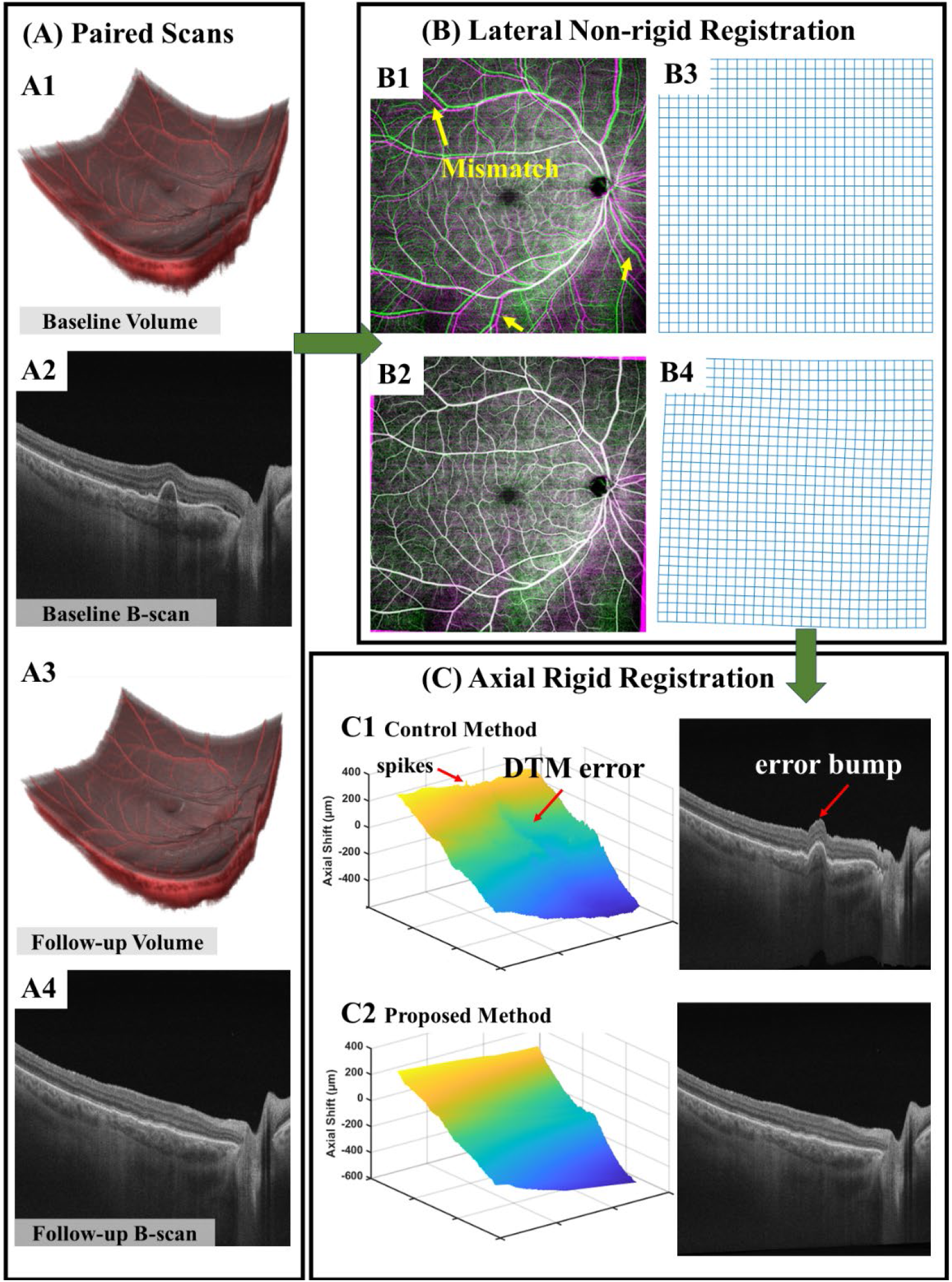
Flowchart of proposed volumetric tracking algorithm. (A) Paired OCT/A volumes and representative B-scans from different visits: the baseline scan serves as the reference volume, and the follow-up scan acts as the moving volume to be registered. (B) Lateral non-rigid registration showing the *en face* angiogram images, projected from the entire depth, before registration (B1) and after registration (B2), as well as the accompanying transformation mesh in initial form (B3) and after lateral registration (B4). Note that the mismatch (yellow arrows) between the scans were corrected based on vasculature pattern. (C) Axial rigid registration. (C1) Depth transformation matrix (DTM) with control method. Note that A-lines are erroneously aligned as error bump in the change region due to DTM errors. (C2) Updated DTM with proposed method. Note that A-lines are appropriately aligned for all regions.

To register the OCT/A volumes in the axial direction, we first selected the pixels with reflectance greater than their mean value across depth profile to avoid intensity artifacts and noise from the background. Then, we calculated the cross-correlation of A-line profiles to determine the depth transformation matrix (DTM), which was obtained from the axial shift components corresponding to the maximum correlation of A-lines [28]. However, treatments have resulted in varied degrees of SRF absorption, leading to reattachment of the retina and resolution of PED (Fig. 1 A2, A4). Our prior algorithm [28] often led to DTM errors (Fig. 1 C1) for those changed regions and consequently the corresponding A-line profiles were erroneously shifted (error bump in Fig. 1 C1). Additionally, local minor mismatches also exist in some A-lines shown as spikes in the DTM. To address this, we refined our algorithm for a more accurate DTM. Firstly, we applied a Gaussian filter to the DTM, subtracted the original DTM to identify the localized mismatch A-lines, and performed interpolation to fix those mismatches. Secondly, since the DTMs are monotonous, we fitted their polynomial trend and then detrended the DTM. For those regions with significant structural change, the DTM has large deviation from its trend and therefore causes a large detrended value. We set a threshold (30 µm) for the detrended matrix to identify those regions and aligned the DTM with its trend in those areas. Subsequently, the updated DTM was employed for axial rigid alignment, resulting in the appropriate shifting of the A-line profiles (Fig. 1 C2).

It should be noted that the proposed volume registration algorithm did not require prior layer segmentation, making the process more efficient and less time-consuming. For assessing the algorithm’s effectiveness in the lateral direction, we obtained the large vessel binary masks from the superficial vascular complexes (SVC). We then computed the Jaccard coefficient, which is defined as the ratio of the intersection to the union of the large vessel binary masks from both baseline and follow-up scans, before and after registration. To evaluate the algorithm’s efficiency in the axial direction, we determined the center of mass (COM) for each A-line post-registration and calculated the differences between the COM maps of the baseline and follow-up scans. Through these quantifications, we were able to evaluate the performance of the proposed registration algorithm both laterally and axially.

### 2.4.3 -D change map

To account for variations in signal strength that may affect OCT reflectance values and to ensure a reliable volumetric comparison after registration, we proposed a compensation method to match the reflectance in baseline and follow-up scans. This was done by 1) generating a structural *en face* image through maximum projection of the OCT signal along the entire A-lines, 2) filling the unregistered edge area in the structural *en face* image using interpolation, 3) applying 2-D Gaussian filters to delineate the trend of localized signal strength but not pick the pattern of blood vessels, and 4) compensating the OCT volume reflectance values by dividing the smoothed structural *en face* image in each depth plane.

We then detected the 3-D change map (*X*_*C*_) by performing voxel-wise comparison of reflectance values in follow-up scan (*X*_*M*_) with the baseline visit scan (*X*_*R*_) using Eq. (1) (Fig. 2). The distribution of voxel value in 3-D change maps from normal control eyes was used to determine the 97.5-percentile cutoff point (0.019) as the contiguity threshold. Voxels in the 3-D change maps (Fig. 2 D-E) with values greater than the threshold were determined as change voxels and further included to calculate the change volume and area.

**Fig. 2.**
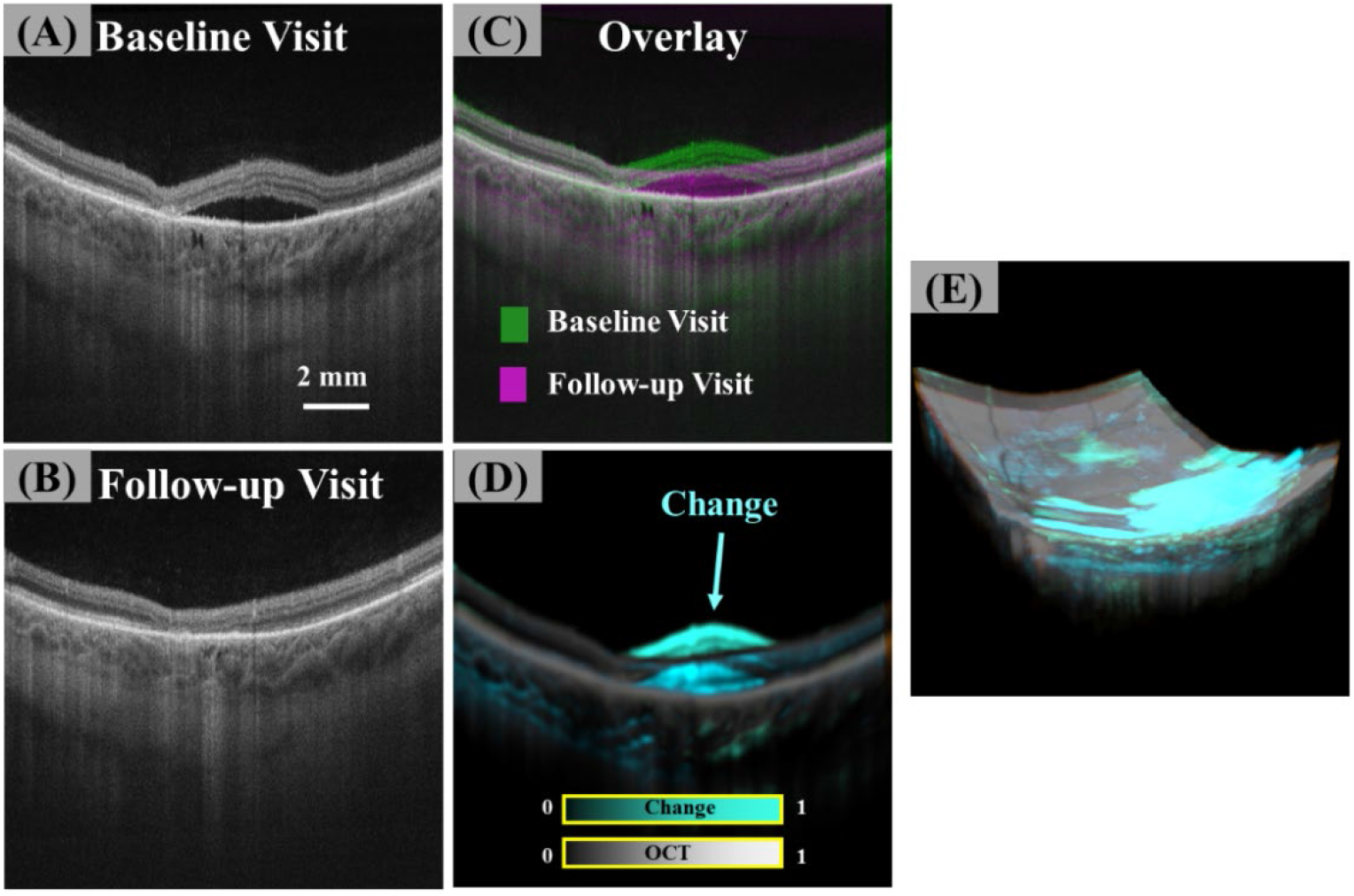
Representative 3-D change map detection for the left eye of a 57-year-old man with central serous chorioretinopathy (CSCR) underwent photodynamic therapy after baseline visit. (A) Baseline visit B-scan displaying accumulated fluid in the subretinal space. (B) B-scan from the follow-up visit showing the resolved fluid. (C) Overlay of the B-scans after registration. (D) Detected 3-D change map (cyan) overlaid with baseline and follow-up B-scans. (E) Detected 3-D change map (cyan) overlaid with baseline and follow-up volumes.

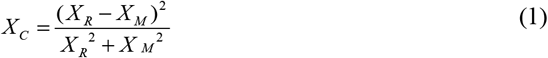

To enable a more refined quantification in the retina and choroid separately, we segmented the inner limiting membrane (ILM) and Bruch’s membrane (BM) layers to differentiate the retinal and choroidal regions. Additionally, 2-D change maps were generated by projecting the 3-D change maps. Jaccard coefficients analysis were applied to the 2-D change maps of the retina and choroid to verify their independence. The unregistered edge area and optic disc-centered 2-mm area were excluded from the analysis. To further examine the components of the 3-D change maps, we focused on the retinal region with graders manually delineated SRF and PED areas on all cross-sectional B-scan images for baseline and follow-up visit and then calculated their changes, and the changes in retinal thickness between visits were calculated by comparing the retinal regions delineated by ILM and BM.

### 2.5 Statistics

Statistical analysis was conducted using SPSS 28.0 (IBM Corporation, Chicago, IL). The Shapiro-Wilk test was performed to assess the normality of quantitative features. Paired-samples Wilcoxon tests were used to compare the SRF and PED between baseline and follow-up visits. Wilcoxon rank-sum tests were used to determine the significance of the 3-D and 2-D structural changes detected between classified study groups. Wilcoxon signed-rank test was used to compare the sum of structural change volume in SRF, PED and retinal thickness, and the change volume calculated from 3-D maps in the retina. The correlation between the sum of structural change volume in SRF, PED and retinal thickness, and the change volume calculated from 3-D maps in the retina was assessed using Pearson correlation analysis. The data were presented as mean ± standard deviation (SD). All P values were 2-sided and considered statistically significant if the value was less than 0.05.

## 3. Results

### 3.1 Volumetric registration

The effectiveness of the volumetric registration algorithm was first verified by evaluating the Jaccard coefficient, which represented the similarity of large vessel binary masks at baseline and follow-up scans (Fig. 3 A-B). In healthy eyes, the Jaccard coefficients increased significantly from 0.25 ± 0.07 to 0.84 ± 0.01 (P<0.001) following registration. The axial difference, quantified by the A-line COM, was negligible at 1.94 ± 2.06 µm for the baseline scans and registered follow-up scans. In CSCR eyes, the Jaccard coefficients were also increased significantly from 0.27 ± 0.11 to 0.83 ± 0.01 (P<0.001) after registration (Fig. 3 C). The difference of COM in patients with CSCR showed strong contrast in the changed regions to consistent regions (Fig. 3 D).

**Fig. 3.**
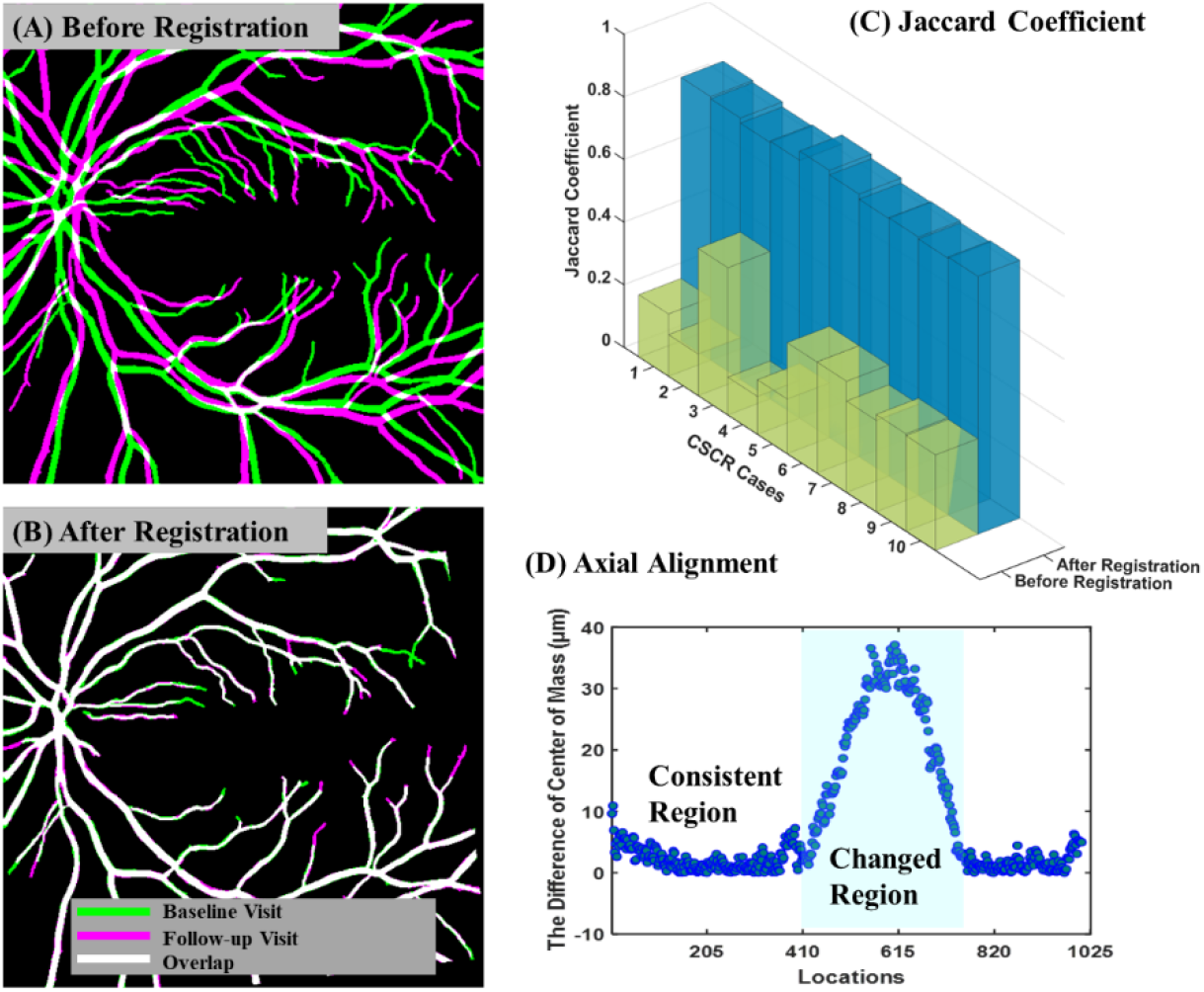
(A) – (B) The large vessel binary masks of superficial vascular complexes (SVC) from baseline and follow-up scans before (A) and after (B) registration. (C) The Jaccard coefficient of the binary large vessel masks was significantly improved, indicating the effectiveness of volumetric registration in the lateral direction. (D) The difference of center of mass (COM) for A-lines in B-scan at Fig. 2 (D) showing strong contrast in the changed region to consistent region.

### 3.2 Structural changes in CSCR eyes

The 3-D structural changes between baseline and follow-up visits were 10.21 ± 6.48 mm^3^ in the treatment group, indicating a considerable morphological cure of the eye following the treatments. Interestingly, the changes were observed not only in the retinal regions (7.79 ± 4.09 mm^3^), but also in the choroidal regions (2.42 ± 2.43 mm^3^). The structural changes in the treatment group were significantly greater than those in the observational group (Total: 0.30 ± 0.42 mm^3^, P<0.01, Retina: 0.21 ± 0.34 mm^3^, P<0.01, Choroid: 0.09 ± 0.12 mm^3^, P=0.03) (Table 1).

**Table 1.** Detected Structural Changes Among Study Groups.

|  | Observational Group | Treatment Group | P-value |
| --- | --- | --- | --- |
| Change volumes in total region ($\text{mm}^3$) | $0.30 \pm 0.42$ | $10.21 \pm 6.48$ | $< 0.01$ |
| Change volumes in retina region ($\text{mm}^3$) | $0.21 \pm 0.34$ | $7.79 \pm 4.09$ | $< 0.01$ |
| Change volumes in choroid region ($\text{mm}^3$) | $0.09 \pm 0.12$ | $2.42 \pm 2.43$ | 0.03 |
| Change area in total region ($\text{mm}^2$) | $4.67 \pm 4.66$ | $85.06 \pm 20.96$ | $< 0.01$ |
| Change area in retinal region ($\text{mm}^2$) | $3.97 \pm 3.96$ | $83.93 \pm 17.12$ | $< 0.01$ |
| Change area in choroid region ($\text{mm}^2$) | $1.36 \pm 1.82$ | $49.18 \pm 19.46$ | 0.02 |
| Change volumes of SRF ($\text{mm}^3$) | $0.01 \pm 0.02$ | $0.92 \pm 1.53$ | 0.19 |
| Change volumes of PED ($\text{mm}^3$) | $0.13 \pm 0.29$ | $0.65 \pm 0.72$ | 0.06 |
| Change volumes of thickness ($\text{mm}^3$) | $0.03 \pm 0.05$ | $4.63 \pm 1.53$ | 0.02 |
| Sum of SRF, PED, Thickness Change ($\text{mm}^3$) | $0.17 \pm 0.35$ | $6.20 \pm 2.79$ | 0.02 |

Similarly, the 2-D *en face* projected structural changes area were significantly greater in the treatment group than in the observational group in all regions measured (Total: 85.06 ± 20.96 mm^2^ vs. 4.67 ± 4.66 mm^2^, P<0.01; Retina: 83.93 ± 17.12 mm^2^ vs. 3.97 ± 3.96 mm^2^, P<0.01; Choroid: 49.18 ± 19.46 mm^2^ vs. 1.36 ± 1.82 mm^2^, P=0.02) (Table 1). Additionally, the projected 2-D change maps of the retina and choroid in CSCR eyes had low spatial correlations with Jaccard coefficient of 0.25 ± 0.23, indicating the detected changes in choroidal regions were not a reflection of the pathologic changes in retina.

### 3.3 Correspond 3D change maps to pathologies

We manually labeled the pathological SRF and PED (Fig. 4 A), as well as delineated the retinal tissue regions, in both baseline and follow-up scans, to investigate the contents incorporated in the automatically detected 3-D change maps. In the observational group, the SRF volume was measured as 0.01 ± 0.01 mm^3^ at the baseline visit and 0.01 ± 0.02 mm^3^ at the follow-up visit, and the PED volume was 0.01 ± 0.02 mm^3^ at the baseline visit and 0.14 ± 0.29 mm^3^ at the follow-up visit. In the treatment group, the SRF volume was measured as 0.92 ± 1.53 mm^3^ at the baseline visit and 0 ± 0 mm^3^ at the follow-up visit, and the PED volume was 0.64 ± 0.72 mm^3^ at the baseline visit and 0.04 ± 0.03 mm^3^ at the follow-up visit. No significant differences were found in SRF volume change (0.01 ± 0.02 mm^3^ vs. 0.92 ± 1.53 mm^3^, P=0.19) or PED volume change (0.13 ± 0.29 mm^3^ vs. 0.65 ± 0.72 mm^3^, P=0.06) between the observation group and treatment group. The structural change volumes of retinal thickness were significantly greater in the treatment group compared to the observational group (4.63 ± 1.53 mm^3^ vs. 0.03 ± 0.05 mm^3^, P=0.02).

**Fig. 4.**
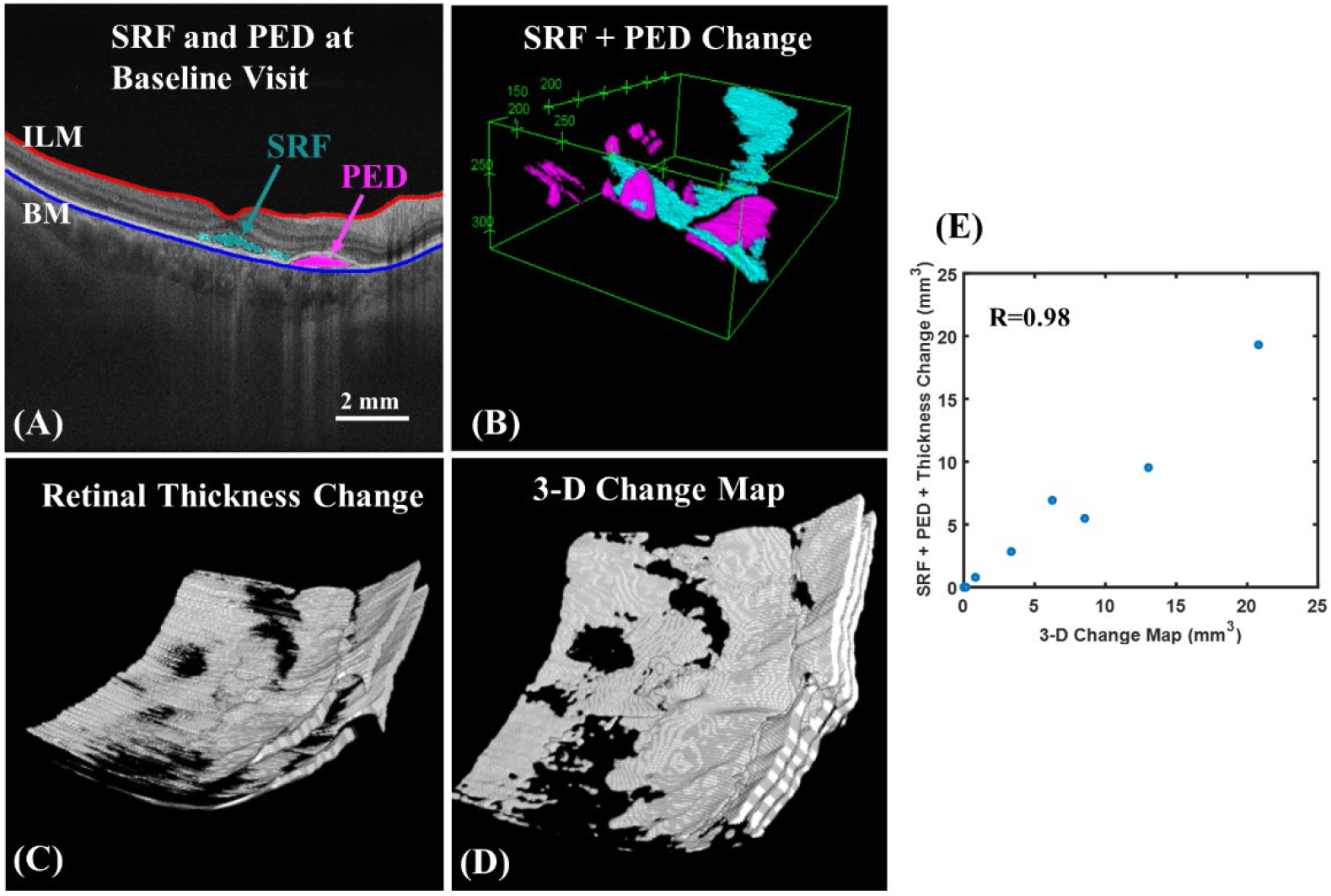
(A) The manually labeled subretinal fluid (SRF) and retinal pigment epithelial detachment (PED) in B-scan image at baseline visit. (B) The SRF and PED change volumes by comparing the labeled binary masks at baseline and follow-up visits. (C) The retinal thickness change volume by comparing the retinal tissue region defined within ILM and BM boundaries at baseline and follow-up visits. (D) The automatically detected 3-D change map. (E) The scatter plot curve of detected 3-D change volume to the sum of SRF, PED and thickness changes.

By summing the manually calculated structural change volumes in SRF, PED and retinal thickness (∑ (SRF + PED + retinal thickness)) in eyes with CSCR, the values were significantly greater in the treatment group compared to the observational group (6.20 ± 2.79 mm^3^ vs. 0.17 ± 0.35 mm^3^, P=0.02). Additionally, the manually calculated change volumes were comparable and showed no significance to the automatically calculated values in the retina region from 3-D change maps in all eyes with CSCR (2.85 ± 3.62 mm^3^ vs. 3.58 ± 4.72 mm^3^, P=0.07), with a high correlation coefficient of 0.98 between the two parameters (Fig. 4 E). It can be concluded that the automatically detected 3-D change map consisted of the pathological SRF or PED, and the resulting retinal thickness change.

## 4. Discussion

This study developed an improved automated volumetric registration algorithm to align OCT/A scans from baseline and follow-up visits in patients with CSCR. The proposed algorithm compensated signal strength variation and can align OCT/A scans regardless of the structural changes. After registration, 3-D structural change maps were obtained to evaluate the progression and resolution of pathologies in CSCR patients by performing volumetric comparison of the scans. We showed that the algorithm achieved great efficiency with the high Jaccard coefficients of large vessel masks laterally and the negligible center of mass of A-lines axially. The structural change in the total, retina, and choroid regions was significantly greater in the treatment group than in the observational group, as evidenced by both 3-D and 2-D measurements.

The axial shift components of the DTM for volumetric registration can potentially introduce errors in structural changes, especially in change areas with SRF absorption and resolution of PED between visits. However, the trends of DTM are monotonous (either progressively increasing or decreasing from one side to the other) and don’t have dips or peaks, therefore, the interpolation and the fitting of trend won’t cause a big problem for the PED regions. Several factors contribute to the monotonous DTM. First, the field of view is the same between the baseline and follow-up scans. Second, the axial length of the eyeball is relatively constant between the baseline and follow-up scans, ensuring the retina maintains its physical shape. Third, any variation in retinal curvature observed in OCT between the baseline and follow-up scans can be attributed to differences in pupil entrance. If the pupil entrance is shifted in a particular direction during the follow-up scan, it will cause an upward curvature on one side and a downward curvature on the opposite side, or vice versa. It is unlike that the different pupil entrance causes dip/peak shape for the DTM, i.e., an upward curvature at both edges of the retina but a downward curvature in the central region. Our results indicated that the structural change was effectively identified in those regions (Fig 2. D). Moreover, the difference in the center of mass prominently distinguished regions of change from consistent regions, and these regions of significant difference align closely with the locations of structural changes observed in B-scans (Fig 3. D). Additionally, the manually calculated change volumes exhibit high congruence with values automatically calculated from the 3-D change maps within the retina region. Therefore, the structural change errors stemming from axial alignment were considered acceptable.

The proposed algorithm has several advantages over traditional method of volumetric registration [31-33]. First, it eliminates the need for prior layer segmentation, which can be a time-consuming and error-prone process. Second, by aligning the entire volume, rather than just specific layers, this algorithm allows for a more comprehensive analysis of the entire retina and choroid with a more accurate overall picture of the disease. It can detect changes that may be invisible in layer-based analysis. Third, although developed and demonstrated in CSCR, the algorithm might be extremely helpful for screening to monitor changes in the retina and choroid over time in health subjects during normal aging to detect early signs of retinal damages. Additionally, the algorithm holds the possibility to be applied in other retinal diseases with more complex structural and vascular changes.

Specifically, we observed that the 2-D *en face* projected structural changes of the retina and choroid were not similar with low Jaccard coefficients, indicating the algorithm was able to capture different patterns of structural change of the retina and choroid over time and calculate them separately. This finding may have important implications for understanding the underlying mechanisms of CSCR progression or conditions as well as for developing targeted and more effective treatments. Moreover, we compared the sum of manually calculated structural change volumes in SRF, PED and retinal thickness, and the automatically calculated change volumes in the retina region from 3-D change maps, the results indicated that the 3-D change maps can reliably detect structural changes of SRF, PED, and thickness caused by CSCR simultaneously.

This study has several limitations. It was retrospective in nature with a small sample size. Despite using FastTrac motion correction, we observed significant motions in certain scans. OCT volumes exhibiting significant motion artifacts were excluded from this study, as these areas could be mistakenly identified as structural changes. Our algorithm was limited by the absence of a more advanced motion correction method. Broader investigations with a more extensive sample size, prolonged follow-up intervals, and consideration of images of varying quality could provide deeper insights. Moreover, the threshold for the detrended DTM to determine the regions with significant structural changes was empirically derived from our current dataset. Further investigation to develop strategies such as adaptive threshold would better adjust the DTM for a more accurate alignment of the A-lines in the baseline and follow up scans to calculate the change. Additionally, the omission of layer segmentation might hinder the appreciation of the changes observed in specific slabs from *en face* images. Lastly, our algorithm operates under the assumption that the pattern of retinal vasculature remains relatively stable over time. As illustrated in Fig. 3B, differences in the vascular binary images at the endpoints of large vessels resulted from segmentation errors and limitation of OCTA rather than changes in the vascular pattern. Nevertheless, in cases with significant vascular dropout or aberrant angiogenesis, the algorithm would require modifications to accommodate these disparities more effectively.

## 5. Conclusion

In conclusion, the study presents an improved automated volumetric registration algorithm designed for aligning OCT scans for comparative analysis. Notably, this algorithm obviates the need for prior layer segmentation and enables the detection of volumetric changes in the entire OCT scan region efficiently. It may show potential to identify the pathology resolution after treatment, as well as tracking recurrence and progression, and can be useful in predicting treatment response and evaluating treatment efficacy in patients with CSCR. Overall, the present study provides valuable insights into the potential use of OCT/A in the diagnosis and management of CSCR and highlights the need for continued research in this area. We hope this algorithm can be integrated into clinical practice to provide a more accurate and efficient way of monitoring the progression of retinal diseases and assessing treatment efficacy in the near future.

## Acknowledgments

We appreciate the funding from the Knight Templar Eye Foundation, Alcon Research Institute, and Eye & Ear Foundation of Pittsburgh to Dr. Shaohua Pi and Dr. Bingjie Wang. We also acknowledge support from NIH CORE Grant P30 EY08098, an unrestricted grant from Research to Prevent Blindness to the Department of Ophthalmology.

## References

1. M. Wang, I. C. Munch, P. W. Hasler, C. Prünte, and M. Larsen, “Central serous chorioretinopathy,” Acta ophthalmologica 86, 126–145 (2008).

2. A. Daruich, A. Matet, A. Dirani, E. Bousquet, M. Zhao, N. Farman, F. Jaisser, and F. Behar-Cohen, “Central serous chorioretinopathy: recent findings and new physiopathology hypothesis,” Progress in retinal and eye research 48, 82–118 (2015).

3. F. Semeraro, F. Morescalchi, A. Russo, E. Gambicorti, A. Pilotto, F. Parmeggiani, S. Bartollino, and C. Costagliola, “Central Serous Chorioretinopathy: Pathogenesis and Management,” Clin Ophthalmol 13, 2341–2352 (2019).

4. D. Petkovsek, and R. P. Singh, “Oral Agents in the Treatment of Central Serous Chorioretinopathy,” in Central Serous Chorioretinopathy (Elsevier, 2019), pp. 293–303.

5. C. Meyerle, “Visual Dysfunction in Central Serous Chorioretinopathy,” in Central Serous Chorioretinopathy(Elsevier, 2019), pp. 21–26.

6. A. Gupta, and K. Tripathy, “Central Serous Chorioretinopathy,” in StatPearls [Internet](StatPearls Publishing, 2022).

7. G. Liew, G. Quin, M. Gillies, and S. Fraser-Bell, “Central serous chorioretinopathy: a review of epidemiology and pathophysiology,” Clinical & Experimental Ophthalmology 41, 201–214 (2013).

8. B. Nicholson, J. Noble, F. Forooghian, and C. Meyerle, “Central serous chorioretinopathy: update on pathophysiology and treatment,” Survey of ophthalmology 58, 103–126 (2013).

9. R. Liegl, and M. W. Ulbig, “Central serous chorioretinopathy,” Ophthalmologica 232, 65–76 (2014).

10. T. J. Van Rijssen, E. H. Van Dijk, S. Yzer, K. Ohno-Matsui, J. E. Keunen, R. O. Schlingemann, S. Sivaprasad, G. Querques, S. M. Downes, and S. Fauser, “Central serous chorioretinopathy: towards an evidence-based treatment guideline,” Progress in Retinal and Eye Research 73, 100770 (2019).

11. M. B. Breukink, A. J. Dingemans, A. I. den Hollander, J. E. Keunen, R. E. MacLaren, S. Fauser, G. Querques, C. B. Hoyng, S. M. Downes, and C. J. Boon, “Chronic central serous chorioretinopathy: long-term follow-up and vision-related quality of life,” Clinical ophthalmology (Auckland, NZ) 11, 39 (2017).

12. D. Huang, E. A. Swanson, C. P. Lin, J. S. Schuman, W. G. Stinson, W. Chang, M. R. Hee, T. Flotte, K. Gregory, and C. A. Puliafito, “Optical coherence tomography,” science 254, 1178–1181 (1991).

13. M. R. Hee, J. A. Izatt, E. A. Swanson, D. Huang, J. S. Schuman, C. P. Lin, C. A. Puliafito, and J. G. Fujimoto, “Optical coherence tomography of the human retina,” Archives of ophthalmology 113, 325–332 (1995).

14. C. A. Puliafito, M. R. Hee, C. P. Lin, E. Reichel, J. S. Schuman, J. S. Duker, J. A. Izatt, E. A. Swanson, and J. G. Fujimoto, “Imaging of macular diseases with optical coherence tomography,” Ophthalmology 102, 217–229 (1995).

15. H. Lee, J. Lee, H. Chung, and H. C. Kim, “Baseline spectral domain optical coherence tomographic hyperreflective foci as a predictor of visual outcome and recurrence for central serous chorioretinopathy,” Retina 36, 1372–1380 (2016).

16. P. Roberts, B. Baumann, J. Lammer, B. Gerendas, J. Kroisamer, W. Bühl, M. Pircher, C. K. Hitzenberger, U. Schmidt-Erfurth, and S. Sacu, “Retinal pigment epithelial features in central serous chorioretinopathy identified by polarization-sensitive optical coherence tomography,” Investigative ophthalmology & visual science 57, 1595–1603 (2016).

17. G. Nkrumah, M. Paez-Escamilla, S. R. Singh, M. A. Rasheed, D. Maltsev, A. Guduru, and J. Chhablani, “Biomarkers for central serous chorioretinopathy,” Ther Adv Ophthalmol 12, 2515841420950846 (2020).

18. I. Maruko, T. Iida, Y. Sugano, A. Ojima, M. Ogasawara, and R. F. Spaide, “Subfoveal choroidal thickness after treatment of central serous chorioretinopathy,” Ophthalmology 117, 1792–1799 (2010).

19. N. Kang, and Y. Kim, “Change in subfoveal choroidal thickness in central serous chorioretinopathy following spontaneous resolution and low-fluence photodynamic therapy,” Eye 27, 387–391 (2013).

20. L. Pertl, A. Haas, S. Hausberger, T. Pichler, D. F. Rabensteiner, G. Seidel, E. M. Malle, and M. Weger, “Change of choroidal volume in untreated central serous chorioretinopathy,” Retina 37, 1792–1796 (2017).

21. E. Costanzo, S. Y. Cohen, A. Miere, G. Querques, V. Capuano, O. Semoun, A. El Ameen, H. Oubraham, and E. H. Souied, “Optical coherence tomography angiography in central serous chorioretinopathy,” Journal of ophthalmology 2015(2015).

22. C. Rochepeau, L. Kodjikian, M.-A. Garcia, C. Coulon, C. Burillon, P. Denis, B. Delaunay, and T. Mathis, “Optical coherence tomography angiography quantitative assessment of choriocapillaris blood flow in central serous chorioretinopathy,” American journal of ophthalmology 194, 26–34 (2018).

23. H. Shiihara, S. Sonoda, H. Terasaki, N. Kakiuchi, T. Yamashita, E. Uchino, F. Murao, H. Sano, Y. Mitamura, and T. Sakamoto, “Quantitative analyses of diameter and running pattern of choroidal vessels in central serous chorioretinopathy by en face images,” Scientific Reports 10, 1–10 (2020).

24. S. Pi, T. T. Hormel, B. Wang, S. T. Bailey, T. S. Hwang, D. Huang, J. C. Morrison, and Y. Jia, “Volume-based, layer-independent, disease-agnostic detection of abnormal retinal reflectivity, nonperfusion, and neovascularization using structural and angiographic OCT,” Biomedical Optics Express 13, 4889–4906 (2022).

25. K. Tsuboi, Y. Guo, J. Wang, E. White, S. Mershon, M. Kamei, D. Huang, Y. Jia, T. S. Hwang, and S. T. Bailey, “Three-dimensional quantification of intraretinal cystoid spaces associated with full-thickness macular hole,” Retina 42, 2267–2275 (2022).

26. E. Borrelli, R. Sacconi, L. Querques, M. Battista, F. Bandello, and G. Querques, “Quantification of diabetic macular ischemia using novel three-dimensional optical coherence tomography angiography metrics,” Journal of Biophotonics 13, e202000152 (2020).

27. B. Wang, A. Camino, S. Pi, Y. Guo, J. Wang, D. Huang, T. S. Hwang, and Y. Jia, “Three-dimensional structural and angiographic evaluation of foveal ischemia in diabetic retinopathy: method and validation,” Biomedical Optics Express 10, 3522–3532 (2019).

28. S. Pi, T. T. Hormel, X. Wei, W. Cepurna, J. C. Morrison, and Y. Jia, “Imaging retinal structures at cellular-level resolution by visible-light optical coherence tomography,” Optics letters 45, 2107–2110 (2020).

29. R. K. Wang, L. An, P. Francis, and D. J. Wilson, “Depth-resolved imaging of capillary networks in retina and choroid using ultrahigh sensitive optical microangiography,” Optics letters 35, 1467–1469 (2010).

30. D. Rueckert, L. I. Sonoda, C. Hayes, D. L. Hill, M. O. Leach, and D. J. Hawkes, “Nonrigid registration using free-form deformations: application to breast MR images,” IEEE transactions on medical imaging 18, 712–721 (1999).

31. M. Niemeijer, M. K. Garvin, K. Lee, B. van Ginneken, M. D. Abràmoff, and M. Sonka, “Registration of 3D spectral OCT volumes using 3D SIFT feature point matching,” in Medical Imaging 2009: Image Processing (SPIE2009), pp. 520–527.

32. H. C. Hendargo, R. Estrada, S. J. Chiu, C. Tomasi, S. Farsiu, and J. A. Izatt, “Automated non-rigid registration and mosaicing for robust imaging of distinct retinal capillary beds using speckle variance optical coherence tomography,” Biomedical optics express 4, 803–821 (2013).

33. P. Zang, G. Liu, M. Zhang, J. Wang, T. S. Hwang, D. J. Wilson, D. Huang, D. Li, and Y. Jia, “Automated three-dimensional registration and volume rebuilding for wide-field angiographic and structural optical coherence tomography,” Journal of biomedical optics 22, 026001–026001 (2017).

